# Isolation of Full Size BAC Inserts by DNA Gap Repair in *E. coli*

**DOI:** 10.1101/549634

**Authors:** George T. Lyozin, Luca Brunelli

**Author notes:** Correspondence should be addressed to L.B.

## Abstract

DNA polymers can comprise millions of base pairs and encode thousands of structural and regulatory genetic elements. Thus, the precise isolation of specific DNA segments is required for accurate gene dissection. Although polymerase chain reaction (PCR) is a standard tool for this purpose, increasing DNA template size leads to the accumulation of polymerase errors, hindering the precise isolation of large-size DNA fragments. Unlike PCR amplification, DNA gap repair (DGR) is a virtually error-free process. However, the maximal size of bacterial artificial chromosome (BAC) insert isolated so far by recombination-mediated genetic engineering (recombineering) is <90 Kilobase pairs (Kbp) in length. Here, we developed a compact bacteriophage P1 artificial chromosome (PAC) vector, and we used it to retrieve a DNA segment of 203 Kbp in length from a human BAC by DGR in *Escherichia coli* (*E. coli*). We analyzed the efficiency of DGR with repressed (recombineering^-^) and derepressed lambda phage *red* genes (recombineering^+^). We showed that both DGR efficiency and the percentage of PAC clones containing the expected 203 Kbp BAC insert improved with increasing size of homology arms. In recombineering^+^ *E. coli* cells and with an efficiency of electroporation of 8×10^9^/1µg pUC plasmid DNA, DGR efficiency and the percentage of correct PAC clones were about 5×10^-6^ and 1% for 30 bp; 6×10^-6^ and 30% for 40 bp; and 1.5×10^-5^ and 80% for 80 bp homology arms, respectively. These data show that using long homology arms and a newly developed vector, we isolated for the first time nearly a full size BAC insert with a frequency of correct clones not previously reported.

## Introduction

Bacterial artificial chromosome (BAC) and bacteriophage P1 artificial chromosome (PAC) cloning vectors can stably propagate DNA of hundreds of Kilobase pairs (Kbp) in length.^1, 2^ This size exceeds the average size of a mammalian gene by approximately ten times. In such a large DNA molecule, all native regulatory gene elements are likely to be present along with the gene. As a result, BAC and PAC are popular in studies of gene expression. However, each BAC and PAC system represents the complex interaction of a few genes and several regulatory elements. In order to identify different genetic elements and their function, it is necessary to isolate specific sequences from the native BAC context and to embed them into experimental genetic models. The isolation of specific DNA sequences, including from BAC and PAC libraries, is limited by the location of a specific nuclease cleavage site flanking the sequence of interest. Alternative, more precise approaches are represented by the polymerase chain reaction (PCR) and reactions based on general DNA recombination such as recombination-mediated genetic engineering (recombineering) using lambda phage genes. These genes are powerful and easy to transfer between different *Escherichia coli* (*E. coli*) strains. *E. coli* cells with general recombination phage transgenes can utilize short homology stretches (AKA, homology arms) of approximately 50 basepairs (bp) to drive DNA exchange.^3, 4^ With DNA replacement, one DNA sequence replaces another DNA sequence in a continuous DNA sequence. Otherwise, the DNA sequence that is being replaced can be interrupted by double-stranded DNA break(s) (DSB), leading to discontinuous DNA participating in DNA exchange (Lyozin GT, Brunelli L. DNA Gap Repair-Mediated Site-Directed Mutagenesis is Different from Mandecki and Recombineering Approaches. BioRxiv 313155 [Preprint]. January 17, 2019. Available from: https://doi.org/10.1101/313155). In the latter, which is called DNA gap repair (DGR), there is an actual DNA sequence filling the gap between the homology arms. When a gap is present in a cloning vector, DGR will allow a new DNA sequence to be pasted between the homology arms of the vector (Fig. 1, panel A). DGR cloning is widely used to add wild type sequences to mouse knockout constructs.^5, 6^ However, there are a few requirements for the vectors used in DGR. First, the preliminary introduction of homology arms into the vector should be easier than DNA cloning itself. Second, the vector should be stable upon induction of *red* genes.^7^ Third, since plasmids are frequent vehicles for lambda/lambdoid transgenes,^3, 8–10^ DNA replication of the vector should be compatible with replication of the plasmids residing in the same *E. coli* cell. The DNA insert capacity of the vector should be comparable to BACs since they are a standard DNA source for subcloning. Considering these requirements, mini-PAC plasmids would represent perfect candidates for vectors in DGR cloning.^11^ Here, we show how a newly generated mini-PAC vectors enables the isolation of a full size BAC insert.

**Fig. 1.**
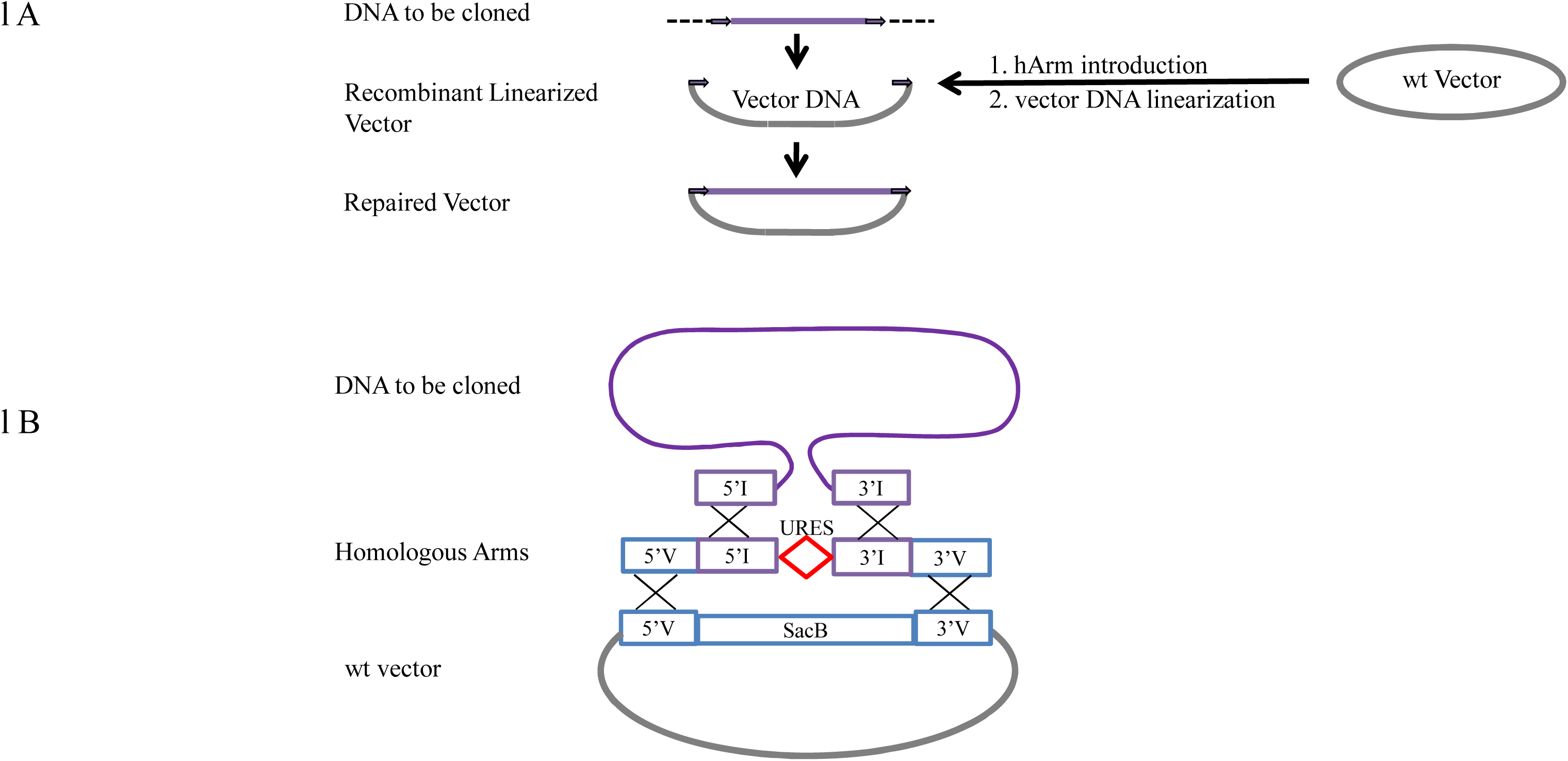
Use of pP1RV2 in DGR cloning. Panel A: representation of DGR cloning. Panel B: structure of the targeting DNA for the introduction of homology arms by recombineering. Arrows (Panel A) and open boxes (Panel B) represent homology arms of a cloning vector and the corresponding homology arms on the DNA to be cloned. 5’V & 3’V: homology arms flanking SacB of the vector; 5’I & 3’I: homology arms to the Insert; URES: a unique restriction endonuclease cleavage site.

## Materials and Methods

### Chemicals and enzymes

Chemicals were purchased from BioBasic (Markham, Ontario, Canada) or ThermoFisher (Waltham, MA). PrimeSTAR GXL, Q5 High-Fidelity and Taq DNA polymerases were purchased from Takara Bio USA (Mountain View, CA), New England BioLabs (Ipswich, Massachusetts) and BioBasic (Markham, Ontario, Canada), correspondingly. Restriction endonucleases were purchased from New England BioLabs.

### Plasmids

Wild-type human *GATA4* BAC, 209.5 Kb RP11-737E8 BAC clone (http://bacpac.chori.org/). pP1RV2, name of the new mini-PAC vector generated in Brunelli’s laboratory.

### DNA isolation and purification

pP1RV2 plasmid DNA was extracted as previously described,^12^ using non-ionic detergent (NID) large scale buffer of the following composition: 1.75 M NH4Cl, 50 mM EDTA, and 0.15% IGEPAL CA-630, RNase A, and lysozyme. BAC plasmid DNA was extracted by alkali.^13^ Native and digested BAC DNA was separated using CHEF mapper (Bio-Rad, Hercules, CA).

### PCR

### Homology arm introduction into wild type pP1RV2

Approximately 8,000 bp wild type DNA vector was amplified via PCR from the primers annealing to SacB flanks and listed in Table 2. PCR mix contained 0.1 μM primers; 0.2 mM dNTP; 5% DMSO, 2.5% glycerol, 20 ng/ml wt vector template DNA and either PrimeSTAR GXL or Q5 High-Fidelity polymerase in the corresponding 1x PCR buffer as recommended by manufactures. PCR included 35 three-step cycles: 96°C for 15”, 57-69°C for 10” and at 69°C for either 9’30” (Q5 High-Fidelity polymerase) or 4’30” (PrimeSTAR GXL polymerase). PCR products were treated with DpnI as previously described,^14^ then precipitated with isopropanol with no additional purification. The DNA pellet was dissolved in electroporation buffer and DNA concentration was measured spectrophotometrically using DS-11 FX+ (DeNovix, Wilmington, Delaware).

### Genotyping bacterial cultures

Every bacterial culture (0.7–1 μl) was transferred into 96-well PCR plates containing 10 μl PCR mix of 1× PCR buffer; 0.2–0.3 μM primers; 0.2 mM dNTP; 0.2 mg/ml BSA (optional); and 10 U/ml Taq DNA polymerase, BioBasic. 5× PCR buffer composition was 100 mM Tris-HCl, pH 8.8, 50 mM (NH_4_)_2_SO_4_, 15 mM MgSO_4_, 0.5% (v/v) Triton X-100, 0.25% (v/v) Tween 20, and 50% (v/v) glycerol. To allow direct gel loading of PCR mixtures, tracking dye, such as 0.001% (w/v) xylene cyanol, were added to the PCR buffer. The PCR program had a 4-min hold at 94-96 °C to lyse bacteria and denature DNA. Annealing, 10–15 s at 55 °C, and extension times were calculated according to a rate of 500 bp/min. The program lasted for 30–35 cycles. Desalted genotyping primers were ordered from Thermo. Long homology arm introducing primers were ordered from Millipore Sigma, PAAG.

### Recombineering

Mini lambda provided by Dr. Shyam K. Sharan (NCI-Frederick) was introduced in DH10B T1 resistant cells, obtained from New England Biolabs (Ipswich, MA) as described previously.^14^

### Bacterial transformation

Electro-competent DH10B T1 phage resistant (New England BioLabs) harboring RP11-737E8 BAC cells were prepared as described previously.^14^ 2 μl (25ng/μl) linear recombinant vector DNA was electroporated at 12,500V/cm at a time constant of 10 msec; for electroporation efficiency of the cells, 5-10 pg of pUC18 plasmid were electroporated at 18,000V/cm at a time constant of 7.5 msec using Gene Pulser Xcell (Bio-Rad, Hercules, CA) into 10 μl competent cells. Bacterial colonies were counted using of Galaxy 330 colony counter (Rocker Scientific, New Taipei City, Taiwan).

## Results

pP1RV2, the mini-PAC vector recently developed in our laboratory,^15^ has a modular structure. Different versions of the vector vary in the counter-selectable markers that they include. One of the versions of the vector is shown in Figure 2. The vector, pP1RV2 (p − for plasmid; P1 − for P1 phage; RV − for retrieving vector; and 2 – the number of the version) has the following useful features. First, counter-selectable *B. subtilis* SacB stuffer flanked with three restriction nuclease sites (multiple cloning site, MCS).^16^ Second, two native P1 phage replicons: P replicon, a lysogenic replicon maintaining the vector as a single copy, and Lytic (L) replicon functioning under lac promotor control. The latter allows the induction of plasmid copy number to a medium-copy-number by isopropyl β-D-1-thiogalactopyranoside (IPTG) which leads to higher yields of DNA for easier DNA isolation.^11^ *sacB* encodes levansucrase, an enzyme catalyzing synthesis of levan, a sugar non-metabolized by *E. coli* cells and whose accumulation is lethal to cells. Due to this gene, only loss-of-function *E. coli* mutants can grow on media containing 2-5% sucrose. This feature can be used for homology arm introductions in two ways. A premade DNA sequence with homology arms separated with a unique nuclease site can be directionally cloned into the vector between the two MCS (Fig. 1, Panel B; see the DNA structure containing 5’I, URES, 3’I). Insertion of homology arms replaces SacB which permits bacteria to grow in sucrose-containing media, i.e., allowing a positive recombinant vector isolation. Alternatively, vector and homology arm DNA (Fig. 1, Panel B; DNA structure containing 5’V, 5’I, URES, 3’I, 3’V) can be co-electroporated into recombineering competent cells. Both vector DNA and recombineering cells can be conveniently premade and used for introduction of all homology arms. In fact, there should be two pairs of homology arms: the external one with homology arms to 5’ and 3’ SacB regions of the vector (Fig. 1, Panel B; 5’V, 3’V) for recombineering with the vector, and the other one - an internal pair for DGR cloning (with homology to the flanks of the DNA to be cloned) (Fig. 1, Panel B; 5’I, 3’I). The vector containing the homology arms for DGR cloning is called a recombinant vector (Fig. 1, Panel A). This DNA needs to be activated for DGR cloning by exposing the homology arms in the ends of the linear vector. Linearization can be obtained by enzymatic digestion of the vector DNA at the unique site separating DGR cloning homology arms.

**Fig. 2.**
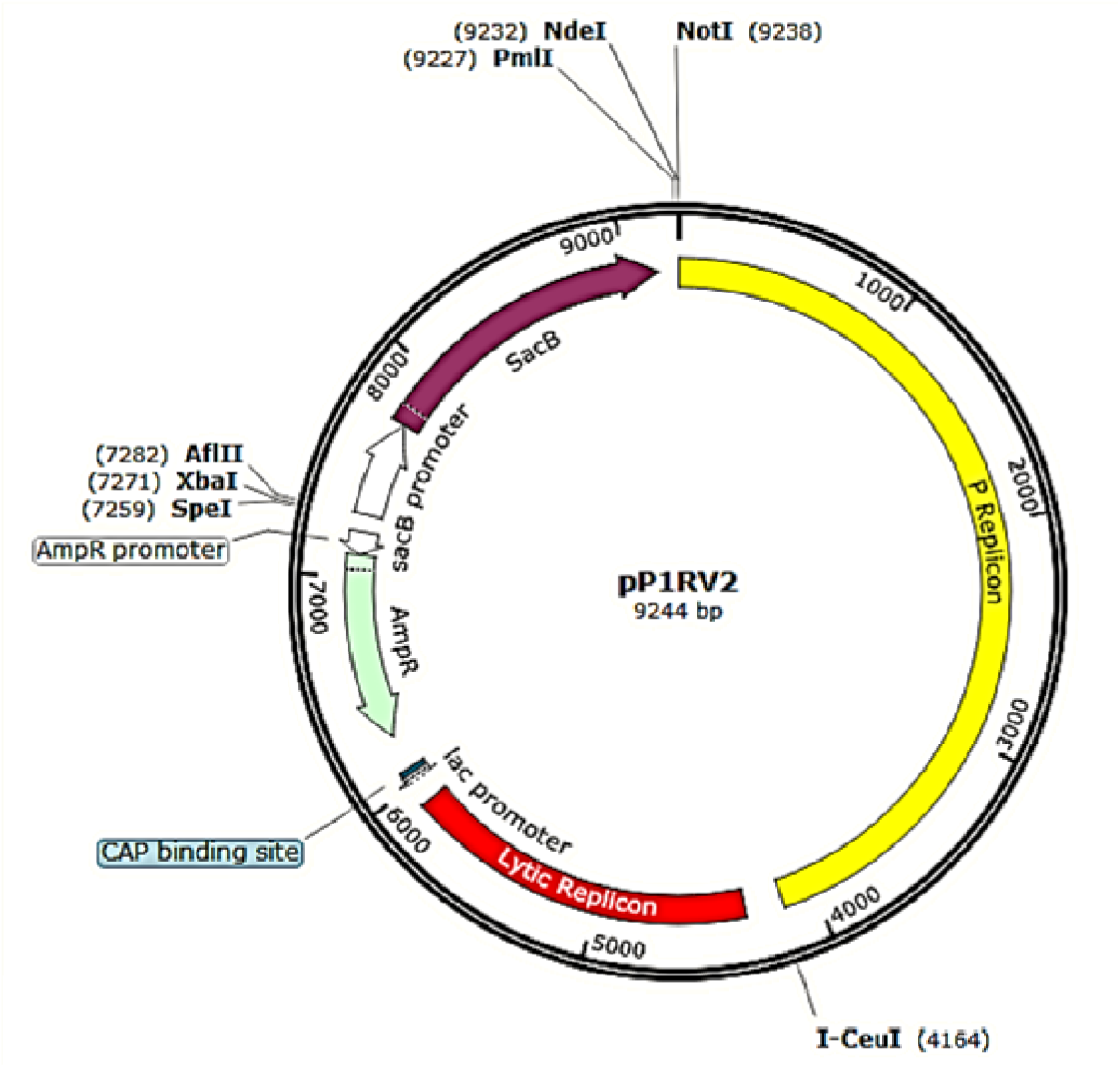
Map of pP1RV2 9.4 Kbp vector version with a counter-selectable Sac B marker

After the electroporation of DNA into cells, a typical by-product of an enzymatic linearization is the background (recombinant circular DNA vector without an insert) cell colonies with a frequency of 10^-4^-10^-5^. (Lyozin GT, Brunelli L. DNA Gap Repair-Mediated Site-Directed Mutagenesis is Different from Mandecki and Recombineering Approaches. BioRxiv 313155 [Preprint]. January 17, 2019. Available from: https://doi.org/10.1101/313155). These background frequencies can make the isolation of repaired cloning vectors containing the insert quite problematic (Fig. 1, Panel A). For example, if 100 cells containing repaired vector are generated per electroporation out of 10^8^ total number of surviving cells, their frequency is 10^-6^. If vector DNA inadvertently produced 10^-4^ background cells, about 10,000 background cells would contaminate 100 cells containing repaired vector.

In this respect, PCR has several beneficial features. First, input DNA template can be both circular and linear, with the latter being unable to establish background colonies. (Lyozin GT, Brunelli L. DNA Gap Repair-Mediated Site-Directed Mutagenesis is Different from Mandecki and Recombineering Approaches. BioRxiv 313155 [Preprint]. January 17, 2019. Available from: https://doi.org/10.1101/313155) Second, the amount of input template is small compared to the amount of output linear PCR DNA products. Third, input DNA template of *in vivo* origin is usually methylated whereas output PCR DNA products are not. Thus, DNA template is sensitive to digestion by nucleases, such as Dpn1, harboring methylated recognition sites. Finally, PCR primers can tolerate inclusion of relatively long non-homologies on their 5’ ends. All of these advantages allow the inclusion of homology arms in the cloning vector and its simultaneous linearization during PCR. Afterwards, digestion of PCR products could remove contaminated wild type DNA template. Therefore, we tested whether PCR can be used for the generation of recombinant vectors from 9.2 Kbp wild type pP1RV2 (Fig. 2 and Fig. 3).

**Fig. 3.**
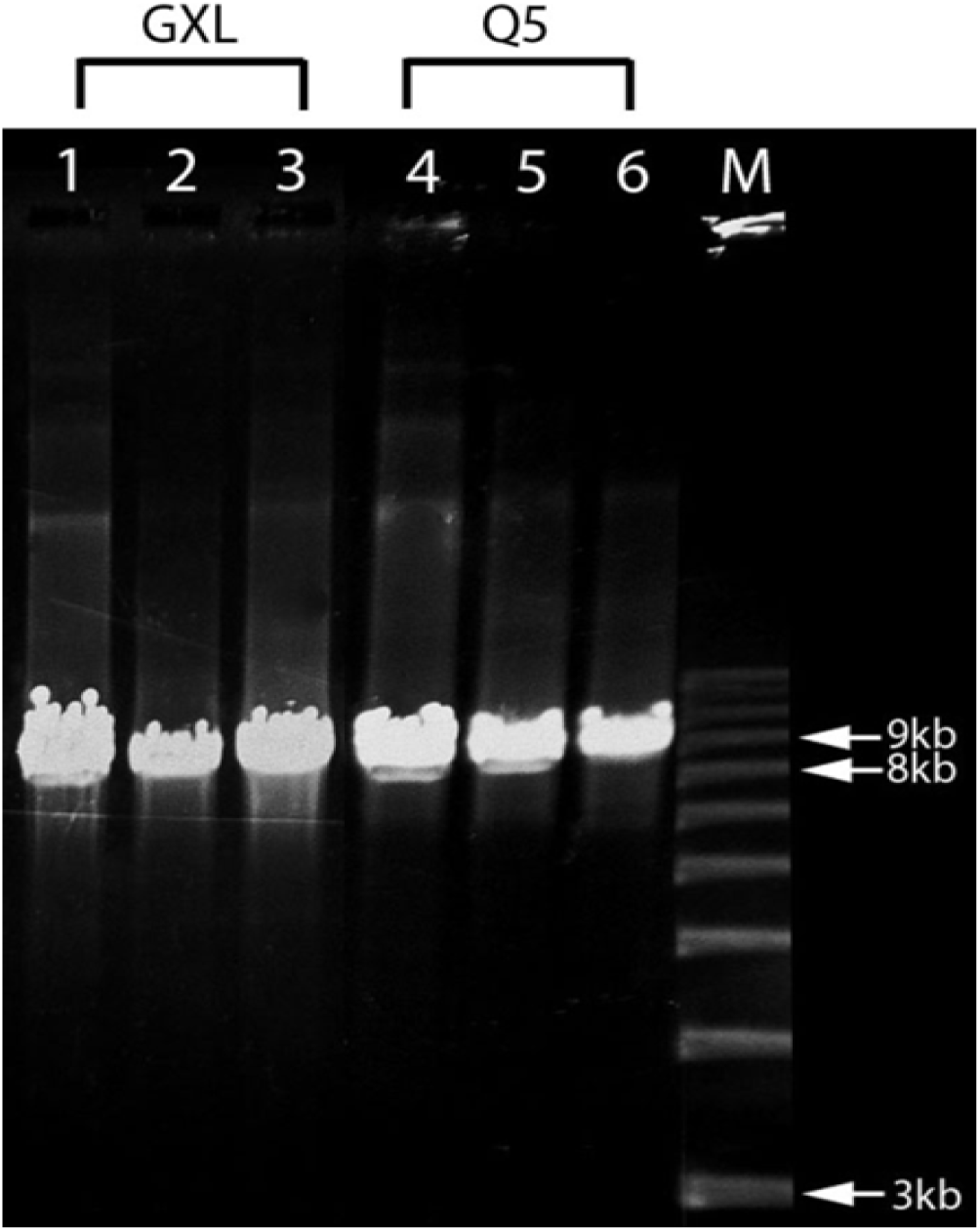
pP1RV2 amplification and insertion of homology arms by PCR. GXL: GXL polymerase; Q5: Q5 polymerase; M: marker; nt: nucleotide. Lanes 1 and 4: PCR with 0.1 µM 20-nt primers complementary to the flanking SacB sequences. 100-nt primers have the same primer binding site as 20-nt primers but with 80 nt non-homology added to 5’ end. Lanes 2 and 5: 0.1 µM. Lanes 3 and 6: 0.5 µM. Primers with up to 100 bp in length can readily amplify ~9 Kbp pP1RV2 allowing simultaneous introduction of homology arms and linearization of the recombinant DNA vector.

In order to amplify ~9 Kbp vector DNA, we tested 5 proofreading polymerases: KOD (Novagen), phusion (Finnenzyme), GXL (Takara), Q5(NEB) and kapa Hi Fi (Kappa Biosystems) with a number of primers of two types, short primers with 100% homology to the template and long primers with ~20 bp homology to the template at 3’ end and non-homologies of various sizes in the 5’-end of the primers. Although we observed some dependency of PCR specificity on the length, sequence and concentration of the primers for all polymerases, we found that the addition of 5% DMSO and 2.5% glycerol as solvent with GXL (Takara), Q5 (NEB) polymerases typically results in a single PCR product of the expected size (Fig. 3, lanes 3, 6). Demonstrated PCR specificity requires no additional DNA purification. Recombinant vector DNA precipitated with isopropanol performed in DGR cloning as well as DNA treated with mung bean nuclease (to remove single stranded DNA) and purified with standard gel electrophoresis (data not shown). These findings allowed us to complete the recombinant vector DNA preparation (including DpnI treatment) within a day.

### Increasing the length of homology arms generates more ampicillin resistant (Amp^r^) colonies in recombineering^+^ cells and less Amp^r^ colonies in recombineering^-^ cells

Wild type pP1RV2 DNA sequence includes more than 30 tetranucleotide DpnI cleavage sites. After treatment with DpnI, PCR-derived recombinant vector DNA can be considered virtually free of *in vivo* wild type DNA template. Linear recombinant vector DNA can be established in DH10B cells only through intramolecular recombination between short terminal sequences resulting in DNA recircularization and establishment of Amp^r^ colonies (Lyozin GT, Brunelli L. DNA Gap Repair-Mediated Site-Directed Mutagenesis is Different from Mandecki and Recombineering Approaches. BioRxiv 313155 [Preprint]. January 17, 2019. Available from: https://doi.org/10.1101/313155). Accordingly, only three types of vectors are expected in DGR cloning: type (1) a vector with no DNA inserts but with an additional microdeletion involving homology arms and adjacent vector sequences, i.e., rearranged recombinant vector; type (2) a vector containing an insert of an unexpected structure and size, i.e., rearranged repaired vector; and type (3) a vector repaired with the correct insert.

Figure 4 demonstrates that in recombineering^+^ cells with increasing homology arm length from 40 to 80 bp, the number of Amp^r^ colonies increased approximately three times. This suggests that increasing homology arm size promotes the generation of type (2) and (3) vectors in which homology is necessary. For recombineering^-^ cells with increasing homology arm length there is a steady decrease of Amp^r^ cell colonies. This suggests that the way in which DNA homologies are used in recombineering^-^ and recombineering^+^ cells is different and is likely connected to a decreasing numbers of colonies in type (1) vector, which does not require any homology between the vector and the BAC. Possible suppression of type (1) vector generation through interaction between the vector and the BAC can be explained by vector DNA entering DGR but, due to the short homology arms, inability of the *E. coli* host to complete the repair process. Since our experiments on DGR of linear pP1RV2 with single-stranded DNA did not support this notion (Lyozin GT, Brunelli L. DNA Gap Repair-Mediated Site-Directed Mutagenesis is Different from Mandecki and Recombineering Approaches. BioRxiv 313155 [Preprint]. January 17, 2019. Available from: https://doi.org/10.1101/313155), we concluded that direct comparison of colony numbers in experiments with electroporation of the same linear recombinant DNA into recombineering^-^ cells with and without homological DNA are needed to clarify this point. In DGR with double-stranded DNA, the colony numbers ratio of recombineering^-^ and recombineering^+^ cells decreased as the size of homology arms increased (Fig. 4). This suggests that the fraction of cells with repaired vectors (vector type 2 and 3) increased. To test this hypothesis, we genotyped bacterial colonies obtained from these experiments.

**Fig. 4.**
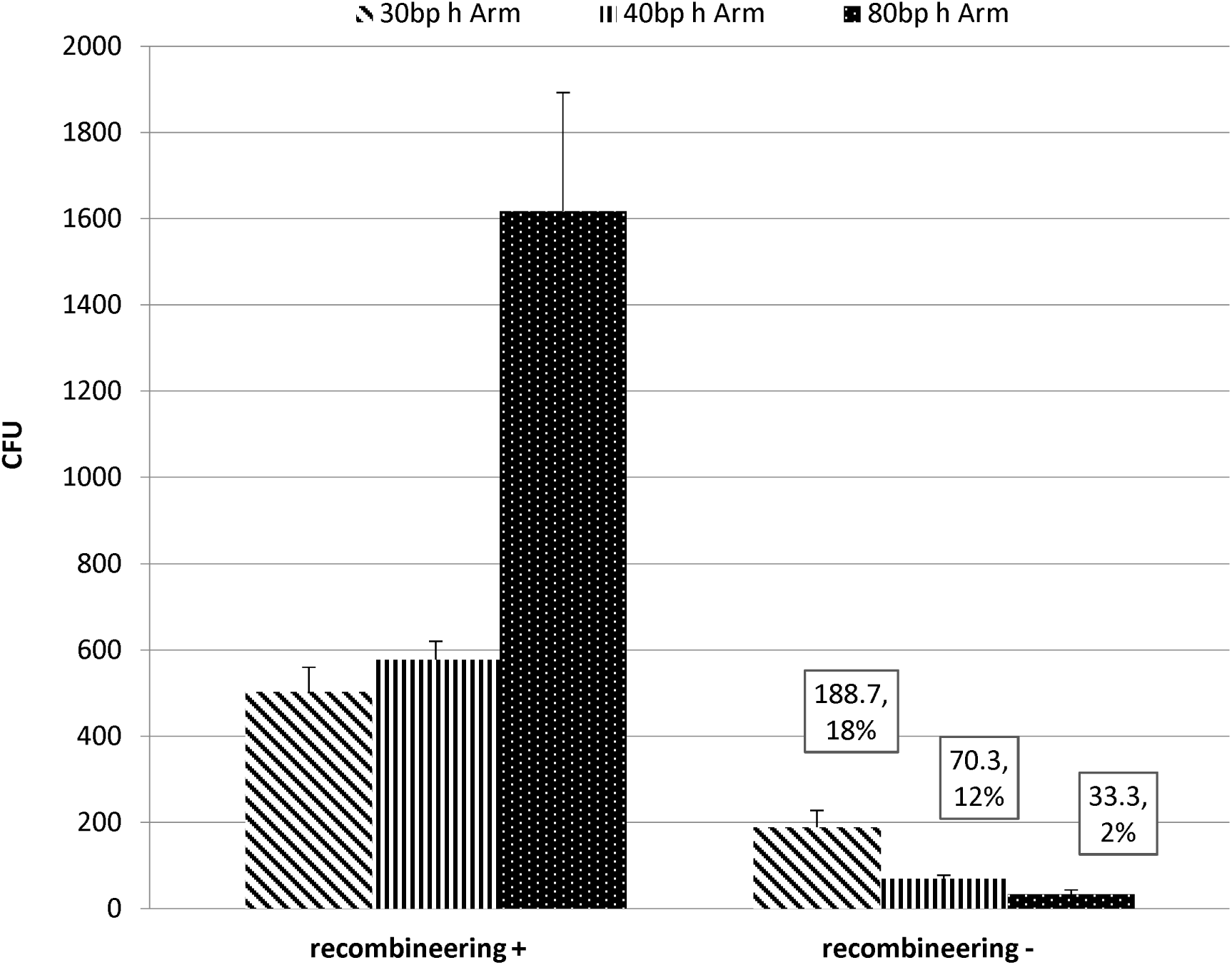
Influence of the size of terminal homology arms of the vector on colony numbers of cells containing the vector repaired by a BAC insert. Linearized recombinant vector DNA (50 ng) with various sizes of homology arms was electroporated into Cm^r^ cells containing a BAC with homological sequences to the homology arms separated by 203 Kbp. The columns with error bars represent average colony numbers per plating. Data labels for recombineering^-^ cells includes the average colony number and its percentage to the corresponding colony number for recombineering^+^ cells

Genotyping bacteria with one primer binding to the vector and the other one binding close to the ends of an insert is useful to detect the physical junction of the vector and the DNA to be cloned.^14^ The method can be applied to bacteria grown in liquid or on solid media. It is both cost and time efficient and particularly suitable for preliminary DNA structure analysis in virtually unlimited bacterial progeny. The genotyping results for both insert ends (break points, BP) are presented in Figure 5 and summarized in Table 1. Application of 30 bp homology arm cloning vectors is not efficient for DGR, with approximately 1% of Amp^r^ BAC insert-containing cells. By extending the homology arm length by 10 bp, the repaired vectors at either or both sides of the insert were detected in approximately 35% of Amp^r^ colonies. The fraction of the insert-containing cells further increased in experiment with 80 bp homology arm cloning vector, reaching about 75%. Since the capacity of PAC vectors exceeds 200 Kbp, and the majority of the colonies were positive for both insert ends junctions with the vector (Table 1), one can expect these colonies to have a full-sized DNA insert without rearrangements (type 3 vectors). To test this hypothesis, plasmid DNA was extracted and analyzed from 16 clones which were PCR positive at both ends of the insert.

**Table 1.**
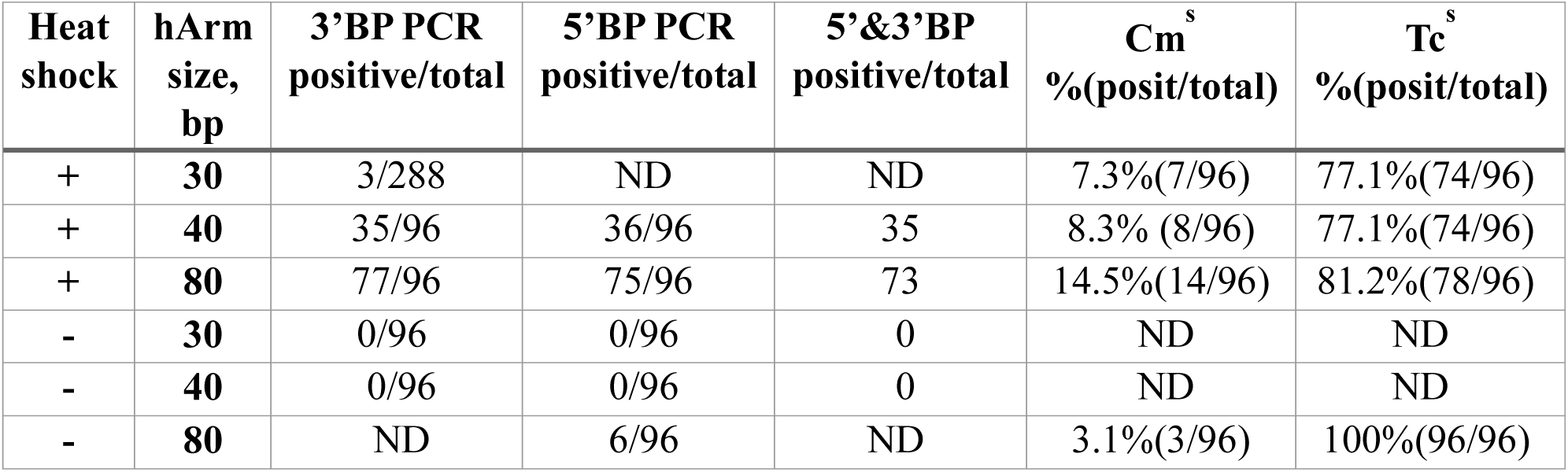
Genotyping and phenotyping results of the colonies from the experiment described in Fig. 4. Genotyping results shown in Fig.5 for + 40 and +80 variants are summarized in the table. Phenotyping was done by transferring the genotyped bacteria from liquid 96 well cultures onto Omnitrays (Nalge Nunc) filled with selective agar. Bacteria were transferred using 96 well replicator (Boekel). **ND**: non determined.

| Heat shock | hArm size, bp | 3'BP PCR positive/total | 5'BP PCR positive/total | 5'&3'BP positive/total | Cm<sup>s</sup> %(posit/total) | Tc<sup>s</sup> %(posit/total) |
| --- | --- | --- | --- | --- | --- | --- |
| + | 30 | 3/288 | ND | ND | 7.3%(7/96) | 77.1%(74/96) |
| + | 40 | 35/96 | 36/96 | 35 | 8.3% (8/96) | 77.1%(74/96) |
| + | 80 | 77/96 | 75/96 | 73 | 14.5%(14/96) | 81.2%(78/96) |
| - | 30 | 0/96 | 0/96 | 0 | ND | ND |
| - | 40 | 0/96 | 0/96 | 0 | ND | ND |
| - | 80 | ND | 6/96 | ND | 3.1%(3/96) | 100%(96/96) |

**Table 2.**
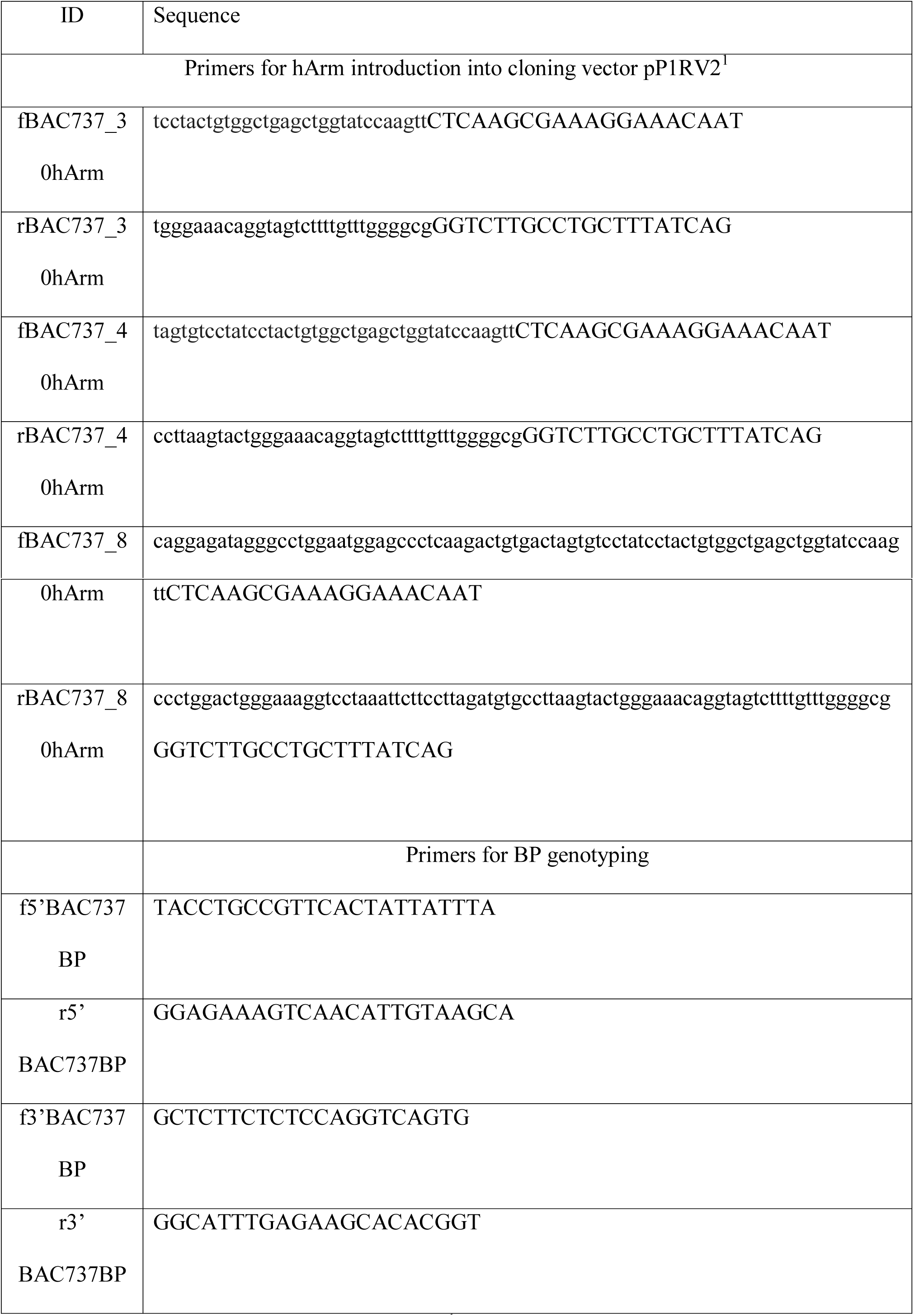
PCR primers used in experiments. ^1^Upper case letters represent primers sequences annealing to vector, lower case letters represent primers sequences homological to the BAC homology arm (30, 40 and 80 bp).

**Fig. 5.**
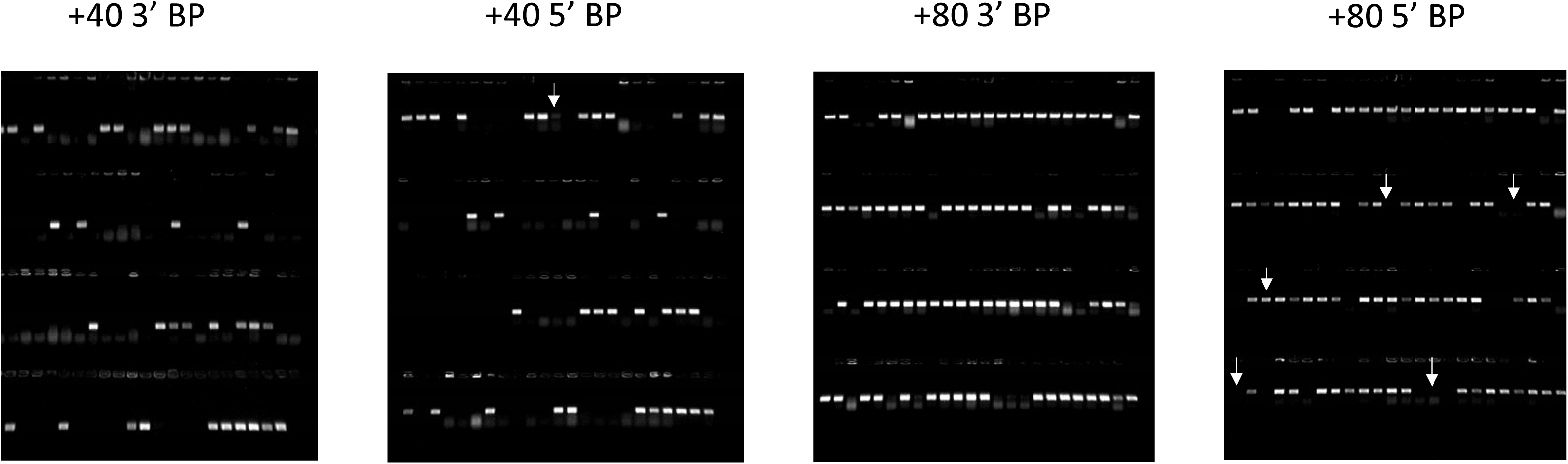
Genotyping bacterial colonies for the physical junction of the insert ends with the vector. Bacteria grown in 96 well plates in liquid media were genotyped using PCR primers annealing close to the insert and vector ends. A 300-400 bp PCR product is expected after insert and vector are joined correctly. +40 and +80 heat colonies were established by heat shocked (recombineering^+^) cells electroporated with the vector DNA including 40 and 80 bp homology arms, respectively. **5’ BP:** 5’ break point (vector-insert junction); **3’ BP:** 3’ break point (insert-vector junction). White arrows represent colonies with a signal only on one end, either 3’ or 5’.

### Isolation of bacterial clones containing repaired vectors with full BAC insert

We utilized mini lambda as a vehicle for *red* genes.^17^ Mini lambda contains genes for site-specific recombination required for lambda phage genome integration/excision in/out of *E. coli* chromosome and resistance to Tetracycline (Tc). Mini lambda genes can be conveniently de-repressed by heat shock. The gene activity can be detected by increasing number of cell colonies with electroporation of a standard targeting vector conferring new drug resistance to the vector component of the plasmid (BAC, PAC) as well as loss of Tc resistance upon mini lambda excision from the *E. coli* chromosome.^14^ Both BAC and PAC vectors are DNA replication compatible. When subcloning a full-size BAC DNA insert into a PAC vector, both plasmids have approximately the same size and can reside in the same cell. To separate the plasmids, plasmid DNA is usually extracted and re-electroporated. Since the chance of BAC and PAC DNA co-electroporation is low, the transformed cells are either Cm^r^, i.e., BAC containing, or Amp^r^, i.e., PAC containing. To skip the plasmid separation step, we phenotyped Amp^r^ colonies for Cm^s^ as well as Tc^s^. The data shown in Table 1 demonstrates that irrespectively of homology arm length about 10% of colonies have lost BAC during DGR, leaving only Amp^r^ vector DNA in the cells. Plasmid DNA was extracted from several Amp^r^/Cm^s^ cells and analyzed with restriction endonucleases by comparing predicted and expected numbers of DNA fragments and their sizes, which confirmed the successful isolation of virtually full-size BAC insert into pP1RV2 (Fig. 6).

**Fig. 6.**
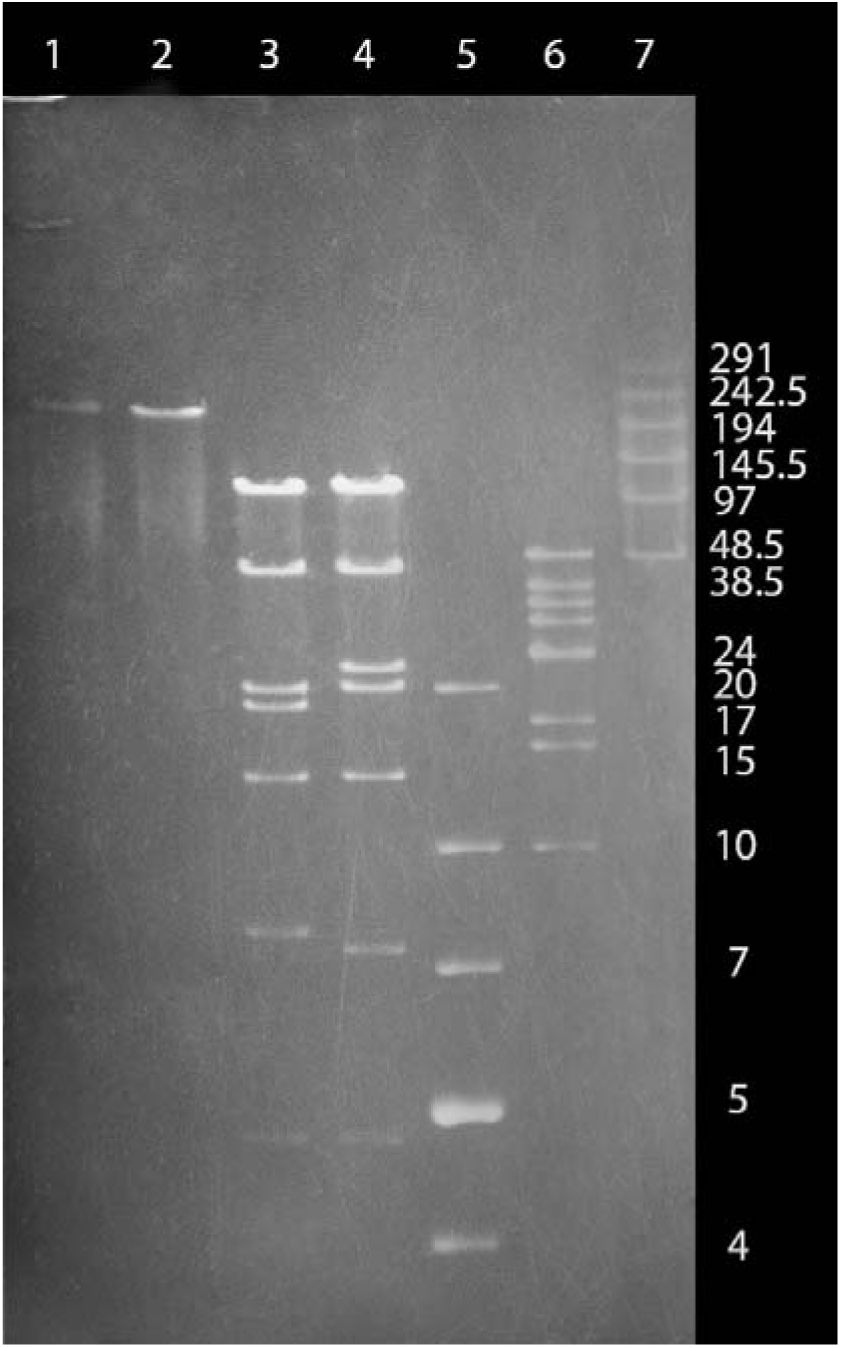
Restriction endonuclease cleavage site analysis of BAC with 209.5 Kbp human *GATA4* region DNA insert and the repaired pP1RV2 containing the 203 Kbp DNA insert isolated from the BAC. Native pP1RV2 was loaded in lane 1. To facilitate DNA molecular size comparison, this DNA was linearized by the homing endonuclease I-CeuI and loaded in lane 2 (note the size is >194 Kbp). The pP1RV2 recombinant DNA (lane 3) and the original BAC DNA (lane 4) digested with NruI and SwaI demonstrate the lack of BAC DNA rearrangements. The expected DNA fragments are as follows: 1) BAC DNA: **4.8 Kbp**, 7 kbp, **13.5 Kbp**, **20.5 Kbp**, 23 Kbp, **43 Kbp**, and **106.5 Kbp**; 2) pP1RV2 DNA: **4.8 Kbp**, 7.8 Kbp, **13.5 Kbp**, **20.5 Kbp**, 18.7 Kbp, **43 Kbp**, and **106.5 Kbp** (the expected common fragments appear in bold). 7 Kbp DNA fragment belongs to the BAC vector DNA, while 7.8 Kbp one represents the pP1RV2 vector DNA (lanes 3 and 4). Lane 5: GeneRuler 1Kb Plus (Thermo-Fisher); Lane 6: lambda DNA-Mono Cut Mix (NEB); Lane 7: concatenated lambda DNA. Pulsed-field gel electrophoresis (5V/cm for 12 hrs.; switch time 2-20 sec.).

## Discussion

These data demonstrate that, by using long homology arms and our newly developed vector, nearly a full size BAC insert with a frequency of correct clones not previously reported can be successfully isolated. Despite these findings, the mechanisms of DGR remain unclear. Two DSBs in circular DNA release two linear DNA fragments (Fig. 7). For simplification, we will designate the smaller fragment an “insert”, and the larger fragment a “vector”. The four ends of the homology arms are numbered from left to right for both “vector” (1v, 2v, 3v, 4v) and insert (1i, 2i, 3i, 4i). In regard to the “vector”, 1v and 4 v ends are internal (ends-in), while as to the “insert”, 1i and 4i ends are external (ends-out).^18^ Based on this difference, one can assume that DNA recombination places DNA with ends-out homology arms into homological DNA, i.e., integrating recombinogenic DNA into homological DNA, whereas DNA recombination places the homological DNA between the ends of the vector with end-in homology arms, i.e., cloning the homological DNA in the vector with recombinogenic ends. These two possible ways of recombinogenic DNA flow (donor or acceptor) are in fact, the two applications of recombineering.^5^ Although there is a strong evidence that the mechanisms of these applications are different,^19^ unlike ends-out recombineering only very limited qualitative data are available regarding ends-in recombineering. In some published data, plasmids with colE1 and pMB1 origins of replication were used as vectors for cloning DNA residing in *E. coli* (chromosomal, and BAC).^20, 21^ These plasmids with 45–52 bp homology arms generated about 400-500 Amp^r^ colonies per electroporation, similar to the colony number provided in Fig. 4 for 40 bp homology arm. In the case of insert sizes of about 25 Kbp, representing the upper limit of the insert in pBluescript plasmid, the fraction of correctly repaired vectors was close to 100%. In pBR322, with a copy number lower than pBluescript, it was possible to clone up to 80 Kbp BAC DNA.

**Fig. 7.**
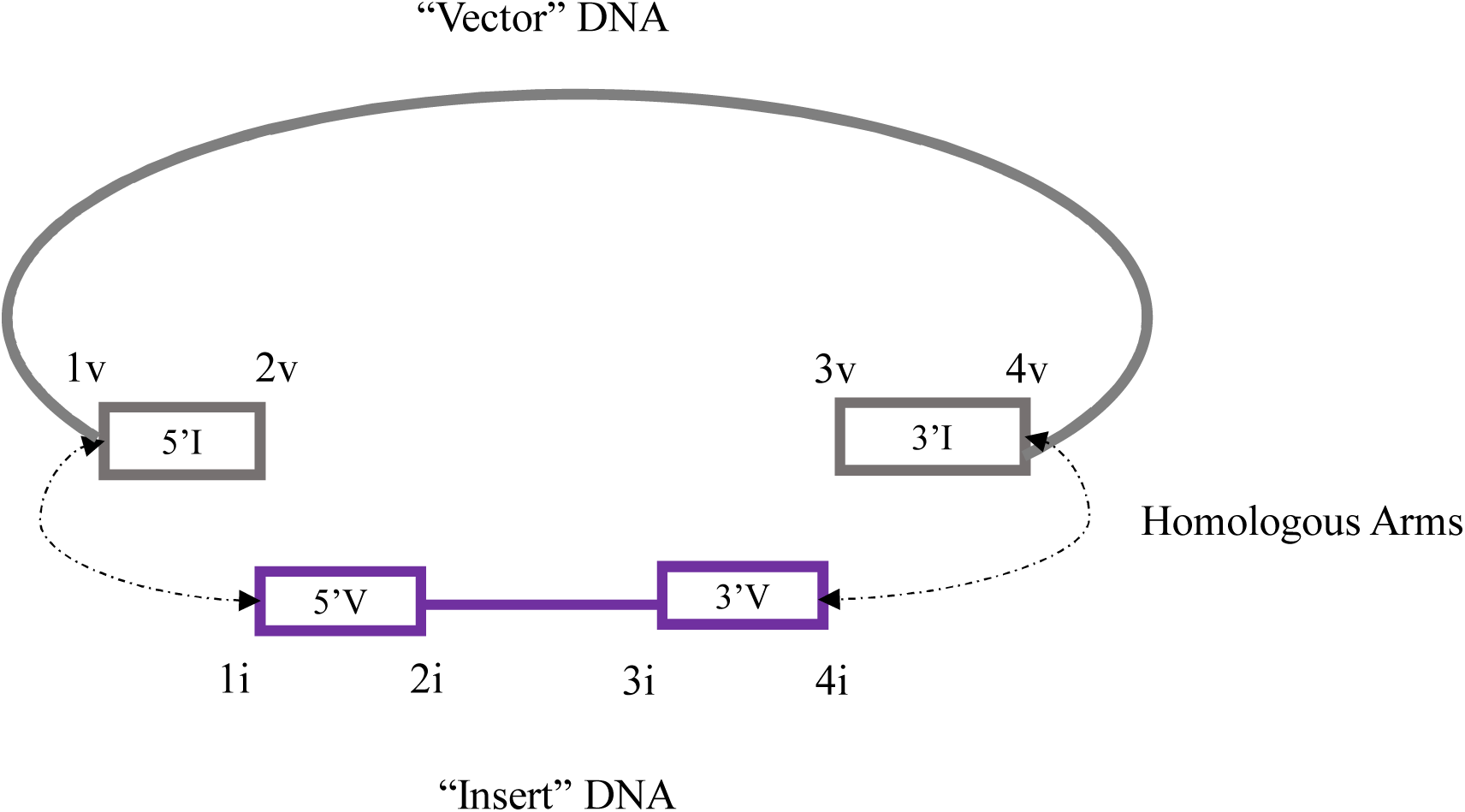
Interpretation of the two possible orientations of the external DNA regions relative to the internal regions. In the “vector” DNA, the internal region is connected *outside* the external DNA regions. Therefore, internal region borders 1 and 4 are internal (ends-in). In the “insert” DNA, the internal region is connected *between* the external DNA regions. As a consequence, internal region borders 1 and 4 are external (ends-out).

However, the fraction of correctly repaired vectors was only about 10%. These findings demonstrate that the problems of DGR cloning are related both to the size of the insert and insert capacity of vector. Since pP1RV2 is a PAC vector, having approximately the same vector capacity as BAC vectors, there is no vector capacity limitation of subcloning BAC DNA into it. Therefore, all problems related to BAC DGR cloning into a pP1RV2 derive only from DGR. The data in Figure 4 indicate that for the full-size BAC insert, increasing the size of homology arms increased DGR precision, which reached about 70% for 80 bp homology arms. For this size of homology arms, the isolation of repaired vectors was possible even from recombineering incompetent cells (Table 1), with about 6% of correctly repaired vectors.

## Conclusion

Here we analyzed intracellular DGR of a linear mini-PAC vector with double-stranded BAC DNA residing in *E. coli* cells. Unlike DGR of the vector with short, single-stranded oligonucleotides, (Lyozin GT, Brunelli L. DNA Gap Repair-Mediated Site-Directed Mutagenesis is Different from Mandecki and Recombineering Approaches. BioRxiv 313155 [Preprint]. January 17, 2019. Available from: https://doi.org/10.1101/313155) this model requires only the intracellular delivery of the DNA vector since homological DNA already resides in *E. coli* cells. With an average of 10^8^ cells surviving electroporation, the frequency of colonies on vector selecting plates were 5×10^-6^ for 30 bp, 6×10^-6^ for 40 bp, and 1.5×10^-5^ for 80 bp homology arms. The fraction of the colonies containing correctly cloned insert DNA was 1%, 30%, and 80%, respectively, Conveniently, the mini-PAC vector used here can be engineered *in vitro* with PCR, it has inducible plasmid copy number, and it is replication-compatible with various plasmids constructed for DNA engineering. As a result, it can be used in different systems of *in vivo* DGR cloning of wild type, mutant, single-stranded, and double-stranded DNA from 21 bp to over 200 Kbp.

## Bibliography

1 Ioannou PA, Amemiya CT, Garnes J, Kroisel PM, Shizuya H, Chen C, et al. A new bacteriophage P1-derived vector for the propagation of large human DNA fragments. Nat Genet. 1994;6(1):84–89.

2 Shizuya H, Birren B, Kim UJ, Mancino V, Slepak T, Tachiiri Y, et al. Cloning and stable maintenance of 300-kilobase-pair fragments of human DNA in Escherichia coli using an F-factor-based vector. Proc Natl Acad Sci U S A. 1992;89(18):8794–8797.

3 Zhang Y, Buchholz F, Muyrers JP, Stewart AF. A new logic for DNA engineering using recombination in Escherichia coli. Nat Genet. 1998;20(2):123–128.

4 Yu D, Ellis HM, Lee EC, Jenkins NA, Copeland NG, Court DL. An efficient recombination system for chromosome engineering in Escherichia coli. Proc Natl Acad Sci U S A. 2000;97(11):5978–5983.

5 Copeland NG, Jenkins NA, Court DL. Recombineering: a powerful new tool for mouse functional genomics. Nature reviews Genetics. 2001;2(10):769–779.

6 Skarnes WC, Rosen B, West AP, Koutsourakis M, Bushell W, Iyer V, et al. A conditional knockout resource for the genome-wide study of mouse gene function. Nature. 2011;474(7351):337–342.

7 Thomason LC, Costantino N, Shaw DV, Court DL. Multicopy plasmid modification with phage lambda Red recombineering. Plasmid. 2007;58(2):148–158.

8 Datsenko KA, Wanner BL. One-step inactivation of chromosomal genes in Escherichia coli K-12 using PCR products. Proc Natl Acad Sci U S A. 2000;97(12):6640–6645.

9 Wu S, Ying G, Wu Q, Capecchi MR. A protocol for constructing gene targeting vectors: generating knockout mice for the cadherin family and beyond. Nat Protoc. 2008;3(6):1056–1076.

10 Datta S, Costantino N, Court DL. A set of recombineering plasmids for gram-negative bacteria. Gene. 2006;379:109–115.

11 Pierce JC, Sauer B, Sternberg N. A positive selection vector for cloning high molecular weight DNA by the bacteriophage P1 system: improved cloning efficacy. Proc Natl Acad Sci U S A. 1992;89(6):2056–2060.

12 Lezin G, Kosaka Y, Yost HJ, Kuehn MR, Brunelli L. A one-step miniprep for the isolation of plasmid DNA and lambda phage particles. PLoS One. 2011;6(8):e23457.

13 Birnboim HC, Doly J. A rapid alkaline extraction procedure for screening recombinant plasmid DNA. Nucleic Acids Res. 1979;7(6):1513–1523.

14 Lyozin GT, Kosaka Y, Bhattacharje G, Yost HJ, Brunelli L. Direct Isolation of Seamless Mutant Bacterial Artificial Chromosomes. Curr Protoc Mol Biol. 2017;118:861–8629.

15 Lyozin GT, Bressloff PC, Kumar A, Kosaka Y, Demarest BL, Yost HJ, et al. Isolation of rare recombinants without using selectable markers for one-step seamless BAC mutagenesis. Nat Methods. 2014;11(9):966–970.

16 Gay P, Le Coq D, Steinmetz M, Ferrari E, Hoch JA. Cloning structural gene sacB, which codes for exoenzyme levansucrase of Bacillus subtilis: expression of the gene in Escherichia coli. J Bacteriol. 1983;153(3):1424–1431.

17 Court DL, Swaminathan S, Yu D, Wilson H, Baker T, Bubunenko M, et al. Mini-lambda: a tractable system for chromosome and BAC engineering. Gene. 2003;315:63–69.

18 Hastings PJ, McGill C, Shafer B, Strathern JN. Ends-in vs. ends-out recombination in yeast. Genetics. 1993;135(4):973–980.

19 Reddy TR, Fevat LM, Munson SE, Stewart AF, Cowley SM. Lambda red mediated gap repair utilizes a novel replicative intermediate in Escherichia coli. PLoS One. 2015;10(3):e0120681.

20 Zhang Y, Muyrers JP, Testa G, Stewart AF. DNA cloning by homologous recombination in Escherichia coli. Nat Biotechnol. 2000;18(12):1314–1317.

21 Lee EC, Yu D, Martinez de Velasco J, Tessarollo L, Swing DA, Court DL, et al. A highly efficient Escherichia coli-based chromosome engineering system adapted for recombinogenic targeting and subcloning of BAC DNA. Genomics. 2001;73(1):56–65.

